# CellTag Indexing: genetic barcode-based sample multiplexing for single-cell genomics

**DOI:** 10.1101/335547

**Authors:** Chuner Guo, Wenjun Kong, Kenji Kamimoto, Guillermo C. Rivera-Gonzalez, Xue Yang, Yuhei Kirita, Samantha A Morris

## Abstract

Single-cell technologies have seen rapid advancements in recent years, presenting new analytical challenges and opportunities. These high-throughput assays increasingly require special consideration in experimental design, sample multiplexing, batch effect removal, and data interpretation. Here, we describe a lentiviral barcode-based multiplexing approach, ‘CellTag Indexing’, where we transduce and label samples that can then be pooled together for downstream experimentation and analysis. By introducing predefined genetic barcodes that are transcribed and readily detected, we can reliably read out sample identity and transcriptional state via single-cell profiling. We validate and demonstrate the utility of CellTag Indexing by sequencing transcriptomes at single-cell resolution using a variety of cell types including mouse pre-B cells, primary mouse embryonic fibroblasts, and human HEK293T cells. A unique feature of CellTag Indexing is that the barcodes are heritable. This enables cell populations to be tagged, pooled and tracked over time within the same experimental replicate, then processed together to minimize unwanted biological and technical variation. We demonstrate this feature of CellTagging in long-term tracking of cell engraftment and differentiation, *in vivo*, in a mouse model of competitive transplant into the large intestine. Together, this presents CellTag Indexing as a broadly applicable genetic multiplexing tool that is complementary with existing single-cell technologies.

## INTRODUCTION

Single-cell technology is advancing at a rapid pace, providing unique opportunities to investigate biological systems and processes with unparalleled resolution. As an increasing variety of assays are being deployed at single-cell resolution, this has presented new challenges for experimental design and data analysis. Recently, batch effects were shown to drive aberrant clustering of the same biological sample processed via two different methodologies^1^, demonstrating how the accuracy of single-cell data analysis can be confounded by measurement errors. Several algorithms currently exist to support the computational correction of batch effects^2–5^. These methods aim to minimize technical artifacts by regressing out known factors of variation during single-cell data processing. However, this requires prior knowledge of the specific factors contributing to batch effects, limiting these approaches. In an alternative strategy, samples are pooled together and subsequently demultiplexed, based on their natural genetic variation^6^, a powerful approach that supports the multiplexing of up to ∼20 samples. However, if the samples are not genetically distinct or are not accompanied by detailed genotypic knowledge, demultiplexing by genetic variation does not represent a feasible approach. For instance, this strategy would not be suitable for comparing different experimental groups from the same individual or animal model where genetic background stays constant.

Recently, several “label-and-pool” approaches have been developed to mark individual cells of the same sample with a distinct barcode prior to pooling and processing in the same single-cell RNA-sequencing (scRNA-seq) run^7–12^. For example, cells can be tagged with barcoded antibodies^9,12^, chemically labeled with DNA oligonucleotides^8,10^, or transiently transfected with DNA oligonucleotides^11^, such that sample identifiers for each cell can be read, in parallel with their transcriptomes. Similarly, several other methods exist to couple genetic perturbations with barcodes^13–17^, although these have not been demonstrated to support reliable, large-scale sample multiplexing. Here, we introduce a methodology to multiplex biological samples via long-term genetic labeling with heritable virally-delivered barcodes, ‘CellTags’. In this approach, defined 8-nucleotide (nt) CellTag barcodes are expressed as polyadenylated transcripts, captured in standard single-cell processing protocols. This design permits the indelible labeling and subsequent identification cells by sample, in parallel with measurement of their identity and state. In contrast to labeling approaches based on transient physical interactions at the cell or nuclear surface, CellTag Indexed cells retain their heritable barcodes for an extended period *in vitro* and *in vivo*, supporting long-term cell tracking experiments. This also distinguishes CellTag Indexing as a unique multiplexing tool in that cell samples can be tagged, mixed and tracked within the same biological replicate, and processed together to mitigate unwanted biological and technical variation.

Here, we validate CellTag Index-based multiplexing via the labeling and mixing of genetically distinct populations, demonstrating accurate and efficient demultiplexing of sample identity. Furthermore, we demonstrate the efficacy of CellTag Indexing for long-term live cell multiplexing, via the establishment of a unique competitive transplant model. In this context, we showcase how CellTag Indexing can be used for *in vivo* multiplexing to precisely quantify engraftment and differentiation potential of distinct, competing cell populations. Together, this positions CellTag Indexing as a broadly applicable tool, easily deployed in cell culture- and transplantation-based assays, that is compatible across different single-cell modalities.

## RESULTS

### Genetic labeling of biological samples via CellTag Indexing

Here, we describe our lentiviral CellTag toolbox for labeling cells with transcribed DNA barcodes, acting as cell/sample identifiers that can be easily recovered from single transcriptomes. CellTag Indexing is based on the integration of defined 8-nt barcodes (CellTags), delivered via lentivirus. In this design, CellTags are positioned in the 3’ UTR of the Green Fluorescent Protein (GFP) gene, followed by an SV40 polyadenylation signal sequence (Fig. 1A). Lentivirus carrying a defined CellTag is used to transduce and genetically label a sample, where GFP is included in this design to enable straightforward quantification of transduction efficiency. This results in the high expression of heritable, polyadenylated CellTag transcripts that are efficiently captured in standard single-cell library preparation pipelines, allowing for the demultiplexing of original sample identity in downstream analysis. We previously demonstrated the efficacy of this approach to label cells with combinations of random CellTags to support lineage tracing in cell fate reprogramming^18^. While this is a powerful approach to track clonally-related cells, it requires more complex experimental design and significant computational analysis. Furthermore, only ∼50% of labeled cells can be tracked via this method; while this supports high-confidence lineage reconstruction, it is not suited to high-efficiency cell labeling for the purpose of sample multiplexing. Our goal here was to expand the utility of CellTagging to support sample multiplexing.

**Fig. 1:**
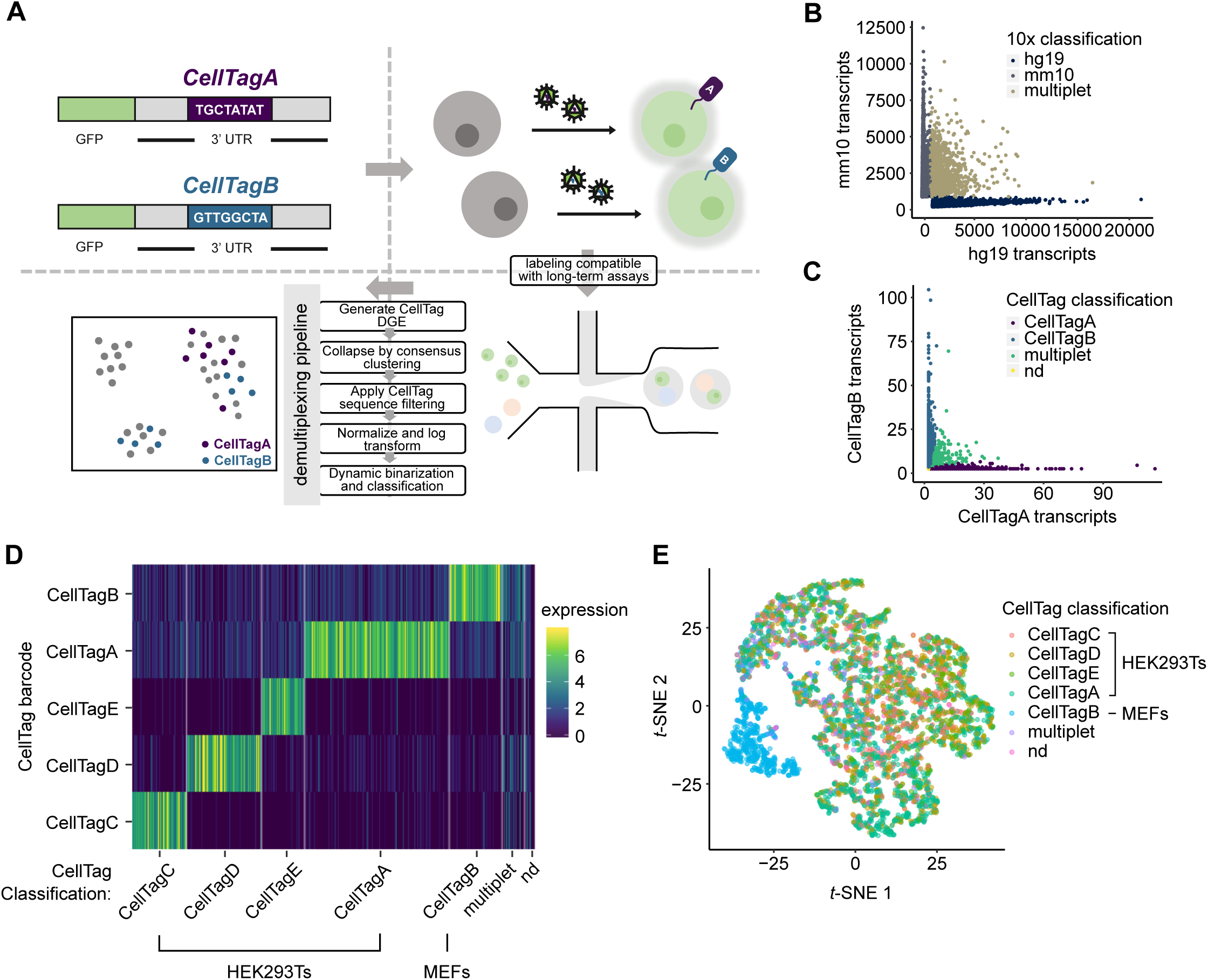
Validation of CellTag Indexing for genetic labeling of biological samples. **A**, Schematic of CellTag Indexing. CellTag barcodes are positioned in the 3’ UTR of a lentiviral GFP construct with a SV40 polyadenylation signal. Barcoded viruses produced from CellTag constructs are used to transduce the cells to be ‘tagged’. Tagged cells can then be pooled for single cell profiling. Prior to analysis, cell identity is demultiplexed by our classifier pipeline: CellTag digital gene expression (DGE) matrix is generated by extracting and counting CellTag sequences for each cell; the DGE is then collapsed by consensus clustering of the detected CellTags; after filtering and log normalization, the DGE is processed by dynamic binarization and classification. Classification results can be visualized as metadata overlaying single transcriptomes projected onto reduced dimensions. **B**, Scatter plot of 18,159 transcriptomes from the 2-tag species mixing experiment, classified by 10x Genomics Cell Ranger pipeline into 9,357 single human cells, 7,456 single mouse cells, and 1,346 multiplets based on alignment to the custom hg19-mm10 reference genome. **C**, Scatter plot of 18,159 transcriptomes from the 2-tag species mixing experiment, demultiplexed by CellTag Indexing into 7,510 human cells (CellTagA), 6,397 mouse cells (CellTagB), 1,040 multiplets, and 3,212 non-determined cells. **D**, Log-normalized CellTag expression of the 4,673 transcriptomes from the 5-tag species mixing experiment, demultiplexed into their respective sample identity on the x-axis; CellTag barcodes, y-axis. **E**, Transcriptomes from the 5-tag species mixing experiment projected onto reduced dimensions by *t*-SNE, visualized with CellTag classification. CellTagC, CellTagD, CellTagE, and CellTagA label HEK293Ts; CellTagB labels MEFs.

First, to ensure that CellTag Indexing does not perturb cell physiology, we tested the impact of labeling on a well-characterized lineage reprogramming system, B cell to induced macrophage reprogramming^19^. We cultured HAFTL pre-B cells and induced reprogramming to macrophage fate with ଲ-estradiol, as previously described^19^. One biological replicate was transduced with CellTag lentivirus, while an independent control replicate, cultured in parallel, was not transduced (Fig. S1A). After 72 hours of reprogramming, the two induced macrophage samples were independently processed for sequencing, along with a sample of the original, untransduced B cells. This yielded 1,310 CellTagged transcriptomes, 2,849 control transcriptomes, and 972 B cell transcriptomes. We detected a median of 6 CellTag transcripts per cell in CellTagged transcriptomes (CellTags were detected in every cell of this sample), and 0 in control transcriptomes (Fig. S1B). Clustering and visualization^5,20^ of CellTagged and control macrophage transcriptomes are interspersed with no independent clustering observed, with both clustering separately from B cells (Figs. S1C&D). Additionally, CellTagged and control induced macrophages exhibit comparable upregulation of macrophage marker expression, accompanied by similar levels of B cell marker downregulation (Figs. S1E). Genome-wide comparison of gene expression of the two samples shows a strong linear association with an R^2^ value of 0.98 (Fig. S1F), confirming that CellTag Indexing does not perturb cell identity or physiology. This is in agreement with our previous study showing that transduction with a random CellTag library does not influence cell behavior^18^.

### Species mixing of genetically distinct cells validates CellTag-based multiplexing

To assess the efficacy of CellTag-based multiplexing, we applied it to ‘species mixing’, an experiment commonly performed to estimate cell co-encapsulation rates in droplet-based scRNA-seq^21^. In this experiment, one sample of human HEK293T cells was labeled with CellTag Index A (CellTagA), and one sample of mouse embryonic fibroblasts (MEFs) was labeled with CellTag Index B (CellTagB), for 24-48 hours. Transduction efficiency was visualized by measuring the percentage of GFP-positive cells (∼90%, Fig. S2B). Labeled cells were pooled, in equal proportions, and processed together for single-cell library preparation and sequencing, yielding a total of 18,159 transcriptomes, with 9,357 single human cells (aligning predominantly to the hg19 genome), 7,456 single mouse cells (aligning predominantly to the mm10 genome), and 1,346 multiplets as classified by 10x Genomics’ Cell Ranger pipeline, based on alignment to the custom hg19-mm10 reference genome (Figs. 1B & S2D). For the purpose of validation, we take this classification result as a benchmark for comparison. To assign sample identity based on CellTag Index expression, we developed a novel demultiplexing algorithm (https://github.com/morris-lab/CellTag-Classifier) that examines the expression distribution of each CellTag Index, followed by a dynamic binarization step to assess each CellTag Index signal on an individual cell basis (Figs. 1A and S2A; see Methods). With this method, we demultiplexed the pooled transcriptomes into 7,510 human cells (CellTagA), 6,397 mouse cells (CellTagB), 1,040 multiplets, and 3,212 non-determined cells (Figs. 1C & S2E). Overall, our algorithm successfully classified, or demultiplexed, 82.3% of all transcriptomes. The presence of non-determined cells is likely due to cells that did not receive sufficient dosage of virus during CellTag Index transduction. This can be enhanced by increasing virus multiplicity of infection (MOI) and visualizing the percentage of GFP-expressing cells prior to sequencing, as demonstrated below. For the purpose of benchmarking, we removed the 3,212 non-determined cells for comparison with the 10x-based classification (Figs. S2C, F, and I). Using Cohen’s kappa as a measure of agreement between independent observations, we calculated a kappa of 0.814 (Fig. S2C), suggesting that our demultiplexing is in strong agreement with the orthogonal 10x-based classification. Furthermore, cells designated as multiplets by both 10x and CellTagging demonstrate a clear increase in the mean numbers of transcripts per cell (Figs. S2G&H), suggesting they do indeed represent multiplets.

To demonstrate the efficacy of CellTag Indexing for multiplexing several biological samples in one experiment, we conducted additional validation where four samples of HEK293Ts and one sample MEFs were transduced with five different predefined CellTag Indexes (HEK293Ts: CellTags C, D, E, and A; MEFs: CellTag B). A total 4,673 cells were sequenced, with an inferred doublet rate of 3.6% (see Methods). Overall, CellTag expression is detected in 99.2% of all cells, reflecting the improved tagging efficiency from an increased MOI. We demultiplexed the transcriptomes as above, including an additional step to resolve misclassified multiplets (Fig. S2A; see Methods). Overall, 4,558 out of 4,673 transcriptomes, or 97.5% of all transcriptomes, were successfully classified (Fig. S3C). Visualization of the classified transcriptomes by heatmap of CellTag barcode expression (Fig. 1D) and by dimension reduction (Figs. 1E, S3A&B) demonstrates clear segregation between species, suggesting that CellTag indexing can be used to reliably multiplex numerous samples.

### CellTag multiplexing enables long-term tracking of cell potential in an *in vivo* competitive transplant model

Current multiplexing methods are based on transient transfection or temporary molecular interactions with the cell or nucleus surface^7–12^. Although relative to CellTag Indexing, this offers faster labeling of cells, it does not support long-term labeling. Here, the unique advantage of CellTag-based multiplexing is that the label is heritable, as a result of stable integration into the cell genome, and can persist for many weeks as we have shown previously^18^. This creates opportunities for the longitudinal analysis of cell behavior over an extended period. Moreover, since experimental groups can be tagged, mixed and tracked within the same biological replicate, unwanted biological and technical variation is minimized. To explore this application of CellTag multiplexing, we applied the method to assess rates of cell engraftment and intestinal differentiation potential in an *in vivo* competitive transplant model.

We previously reported that MEFs can be directly reprogrammed, via forced expression of transcription factors, into progenitor-like cells that possess both hepatic and intestinal potential^22,23^. We demonstrated that these cells, named induced endoderm progenitors (iEPs), are able to functionally engraft a mouse model of induced colitis^23^. Prior to transplant, iEPs possess weak hepatic and intestinal identity, still partially resembling the fibroblasts they originated from. 12-days after transplant into the mouse large intestine, iEPs more closely resemble differentiated intestine^23^. However, in this study, cell identity was assessed via bulk expression analysis that cannot distinguish between different intestinal cell types. Therefore, the mechanism of engraftment and differentiation potential of cells reprogrammed to iEPs remained to be characterized.

Our recent single-cell lineage tracing of fibroblast to iEP reprogramming revealed that this lineage conversion comprises two distinct trajectories: one path successfully reprogramming to iEPs, and an alternate path characterized by progression into a ‘dead-end’ state, where fibroblast identity is re-established^18^. Transition along the successful reprogramming trajectory is accompanied by upregulation of genes such as *Apoa1* and *Cdh1* (E-cadherin). We hypothesized that the Apoa1^High^Ecad^High^ iEP cells constitute the subpopulation responsible for our previously observed colon engraftment^23^. In this context, CellTag Indexing is well-suited for tracking and quantifying reprogrammed and dead-end cell differentiation potential as the barcodes are stably integrated and heritable, making it possible to label cells for long-term tracing transplantation experiments.

To test our hypothesis that the Apoa1^High^Ecad^High^ iEP subpopulation harbors intestinal engraftment and differentiation potential, we first enriched Ecad^High^ and Ecad^Low^ iEP populations using fluorescence activated cell sorting (FACS). Functional assays confirmed that Ecad^High^ iEPs express significantly higher levels of *Apoa1* and *Cdh1*, form larger colonies of reprogrammed iEPs in culture, and retain their Ecad^High^ phenotype, relative to their Ecad^Low^ counterparts (Fig. S4A-C). We then labeled sorted Ecad^High^ iEPs with CellTagA and Ecad^Low^ iEPs with CellTagB, followed by pooling in equal proportions and transplant into a modified mouse model of colonic mucosal injury^24^ (Fig. 2A). Seven days following transplantation, mice were euthanized and dissected, and the engrafted colons collected for histology and single-nucleus RNA-sequencing. Microscopic examination of the engrafted tissue reveals iEP engraftment in discrete patches, located by their GFP expression (Fig. 2B). Histology of the cryosectioned engrafted colon shows the expected tissue architecture with evidence of epithelial injury (Fig. 2C), occasional submucosal iEPs (Fig. 2D), and occasional aggregates of iEPs sitting atop of the damaged epithelium (Fig. S4D).

**Fig. 2:**
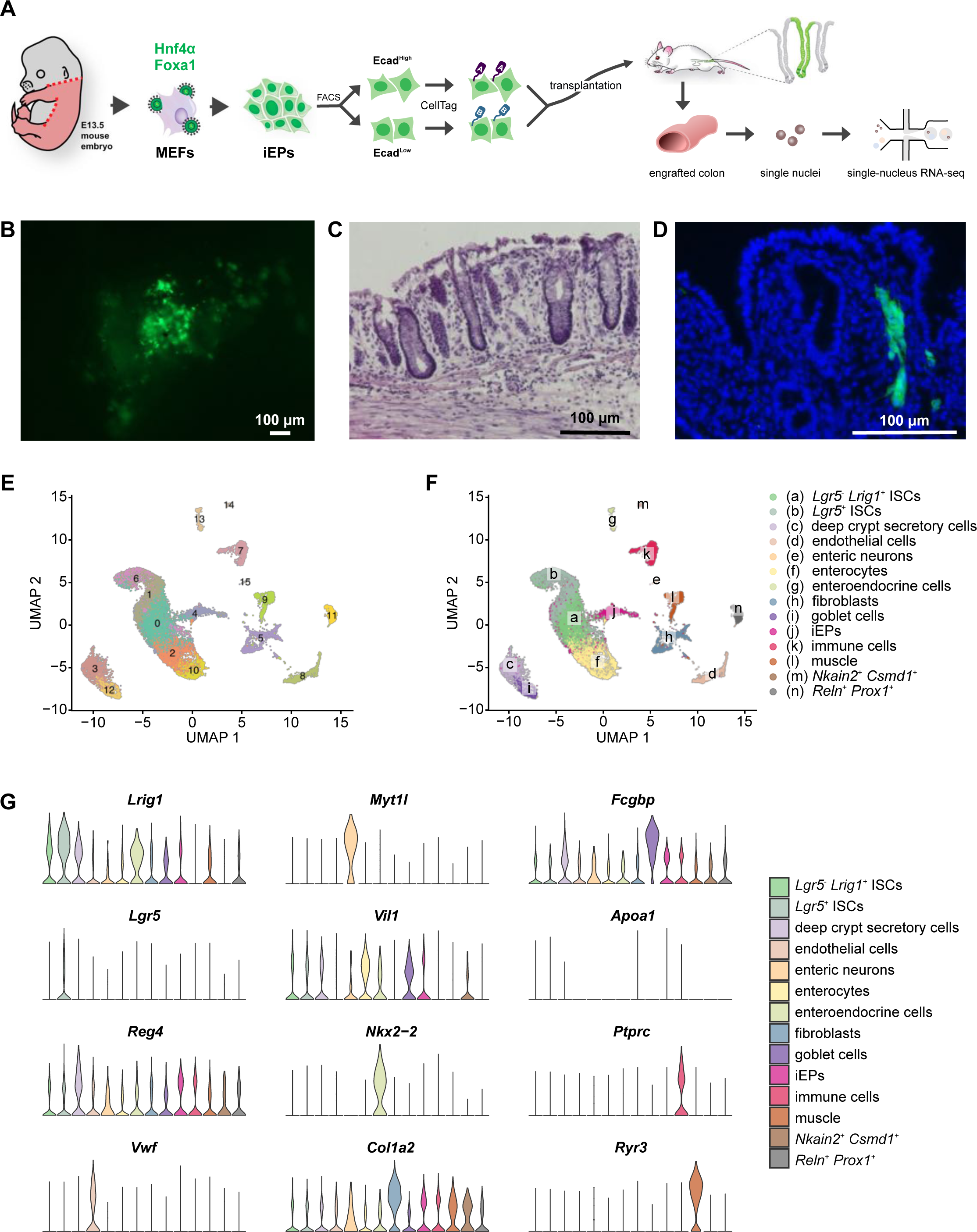
CellTag Indexing for long-term tracking of cells demonstrated in a competitive transplant experiment. **A**, Schematic of iEP generation and enriched into Ecad^High^ and Ecad^Low^ populations by FACS, labeled with CellTagA and CellTagB respectively, pooled in equal proportions and transplanted into a mouse model of colonic injury. Engrafted colon is then processed for single-nucleus RNA-seq. **B**, Fluorescent microscopic images of the lumen of the engrafted colon, showing patches of GFP^+^ iEPs. Scale bar, 100 μm. **C**, H&E stained section of engrafted colon showing normal intestinal architecture with evidence of epithelial injury. Scale bar, 100 μm. **D**, DAPI stained section of engrafted colon showing GFP^+^ iEPs in the mucosa. Scale bar, 100 μm. **E**, Transcriptomes from three post-engraftment colon tissues sequenced and analyzed, visualized by UMAP, revealing 16 clusters. **F**, Annotation of the 16 clusters into (a) *Lgr5*^-^ *Lrig1*^+^ intestinal stem cells (ISCs), (b) *Lgr5*^+^ ISCs, (c) deep crypt secretory cells, (d) endothelial cells, (e) enteric neurons, (f) enterocytes, (g) enteroendocrine cells, (h) fibroblasts, (i) goblet cells, (j) iEPs, (k) immune cells, (i) muscle, (m) *Nkain2*^+^ *Csmd1*^+^ cells, and (n) *Reln*^+^ *Prox1*^+^ cells. **G**, Marker expression in annotated cell types.

Most intestinal cell recovery protocols focus on harvest of the epithelium, neglecting many other cell types that constitute the intestine. Given the range of engraftment phenotypes observed in our above histology analyses, we considered that iEPs may also differentiate toward non-epithelial cell types. Thus, to capture the full spectrum of intestinal cell identities, we opted to use whole tissue single-nucleus extraction, over epithelial isolation and digestion, to process the engrafted colon for RNA-sequencing. Indeed, single-nucleus RNA-sequencing (snRNA-seq) from three colon samples, post-engraftment, followed by Uniform Manifold Approximation and Projection (UMAP)-based visualization^20^ revealed 16 clusters (Fig. 2E), corresponding to a range of different intestinal epithelial cell types. This included intestinal stem cells (ISCs), enterocytes, deep crypt secretory cells, goblet cells, enteroendocrine cells, as well as non-epithelial cell types (endothelial cells, muscle, enteric neurons, immune cells, fibroblasts) (Fig. 2F). To our knowledge, this is the first dataset of such that profiles large intestinal cell types beyond the epithelium. Known intestinal markers are observed such as *Lgr5, Lrig1*, and *Smoc2* in ISCs^25–28^, *Reg4* in deep crypt secretory cells^29^, *Myt1l, Asic2,* and *Syt1* in enteric neurons^30–33^, *Vil1, Plac8*, and *Krt20* in enterocytes^34–36^, *Nkx2-2, Chga,* and *Tph1* in enteroendocrine cells^37,38^, and *Fcgbp, Muc2,* and *Clca1* in goblet cells^39,40^ (Figs. 2G and S5C-E).

Upon further analysis and literature review, we annotated the ISCs into two populations, *Lgr5*^*+*^ ISCs (clusters 1 and 6) and *Lgr5*^*-*^*Lrig1*^*+*^ ISCs (cluster 0), based on distinct patterns of marker expression (Fig. S5C). *Lrig1*, a transmembrane negative regulator of ErbB signaling^41^, is purported to mark a class of ISCs that are phenotypically distinct from *Lgr5*^*+*^ stem cells in the intestine^26,27^, with additional roles in stem cells of the gastric epithelium^42^ and the epidermis^43–45^. *Lgr5*^*+*^ ISCs, located in clusters 1 and 6 in this dataset, express high levels of established intestinal stem cell markers *Lgr5* and *Smoc2*, as well as *Lrig1* (Figs. 2G and S5C-E). In contrast to *Lgr5, Lrig1* is more widely expressed, with moderate levels of expression extending into cluster 0, where *Lgr5* expression is absent (Fig. S5C). This is consistent with two independent studies in the small intestine and colon, where *Lrig1* was expressed in many crypt cells, while the highest levels of *Lrig1* expression were observed in *Lgr5*^*+*^ stem cells^26,27^. Loss of Lrig1 caused crypt expansion in Lrig1-knockout animals, and three-dimensional intestinal spheres derived from Lrig1-knockout animals matured into budding organoids in culture without exogenous ErbB ligands in contrast to wild-type samples^26^. Intriguingly, *Lrig1* was shown to mark a population of ISCs that expand and repopulate the colonic crypt upon tissue damage^27^, although a distinction was not made regarding whether this could be due to the subpopulation of *Lrig1*^*+*^ cells that are also *Lgr5*^*+*^.

Of note, two clusters that remain unannotated (cluster 11, enriched for *Reln* and *Prox1*; cluster 14, enriched for *Nkain2* and *Csmd1*) may represent rare or previously unidentified cell types (Figs. S5D). For example, *Reln* and *Prox1* are known for their roles in neuronal migration^46,47^ and neurogenesis^48,49^; we, therefore, speculate that they may mark a peripheral neuronal cell type in cluster 11.

### Colon engrafted iEPs transition through an intestinal stem cell state

To identify iEPs within the single nucleus landscape of the engrafted colon, we extracted and processed CellTag Indexes across all single transcriptomes. Both CellTagA and CellTagB barcodes were detected in all three post-engraftment samples (Fig. S6A), with clear expression differences between tags (Fig. S6B). Projecting CellTagged iEPs onto the UMAP plot revealed their enrichment in cluster 4 (Figs. 3A&B), while a moderate number of CellTagged cells are found in intestinal epithelial clusters such as cluster 0 (*Lgr5*^*-*^*Lrig1*^*+*^ ISCs) and cluster 1 (*Lgr5*^*+*^ ISCs), expressing ISC markers *Lgr5* and *Lrig1* (Figs. 3C&D, S6C).

**Fig. 3:**
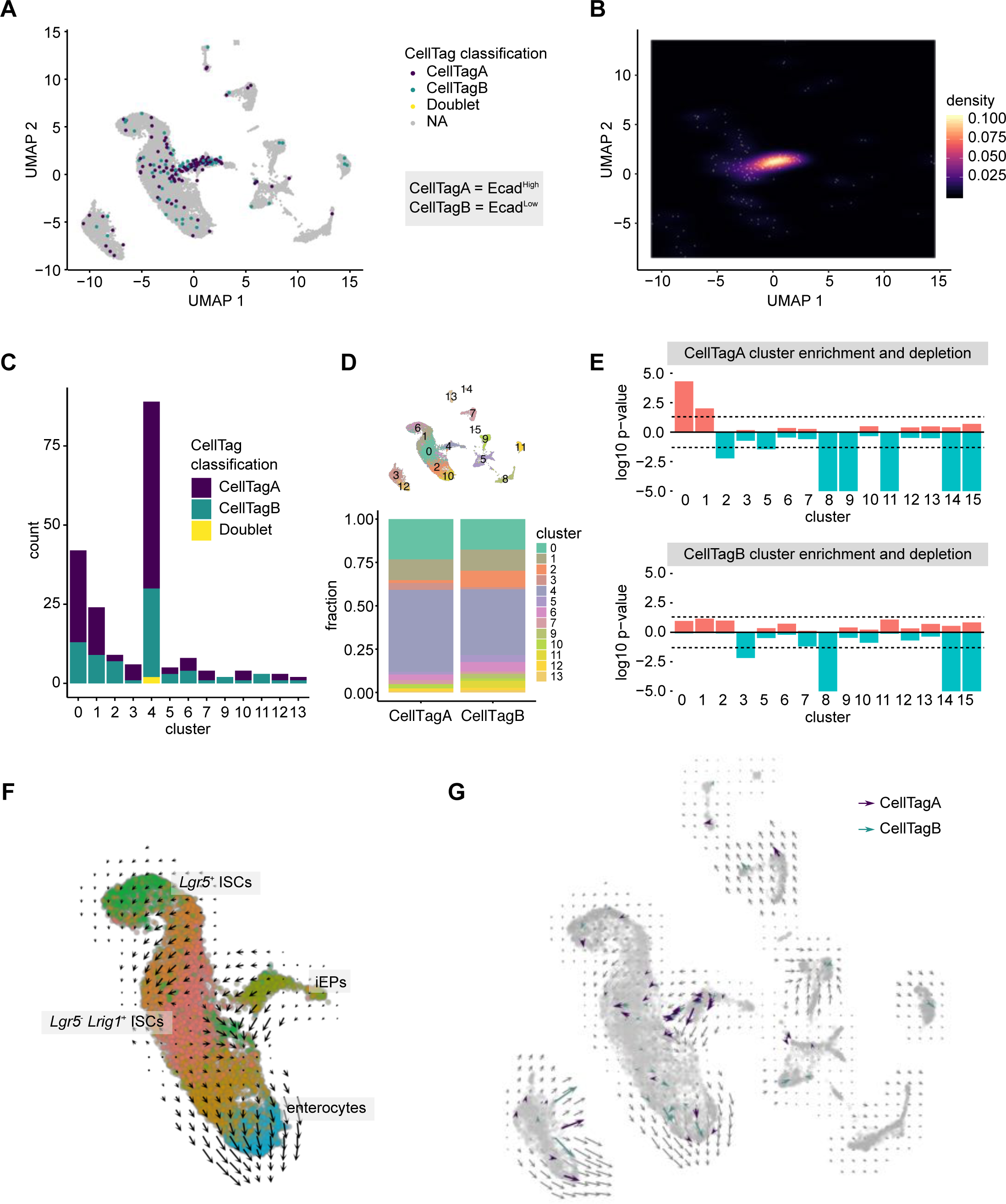
CellTag Indexing revealed iEP engraftment and transition through an intestinal stem cell fate. **A**, CellTags identified engrafted iEPs enriched in cluster 4 (early engraftment iEPs) and the main intestinal epithelial clusters. **B**, Density heatmap confirms enrichment of CellTagged cells in the early engraftment iEP cluster and the main intestinal epithelial cell clusters. **C&D**, Stacked bar plots of CellTagged cells show enrichment in clusters 0, 1, and 4. **E**, Permutation test of cluster enrichment or depletion for each CellTag in intestinal clusters show statistically significant enrichment of Ecad^High^/CellTagA cells in cluster 0 (Lgr5^-^ Lrig1^+^ ISCs, p = 4.03 × 10^-5^) and cluster 1 (Lgr5^+^ ISCs, p = 9.83 × 10^-3^). Y-axis, negative log10 of p-value for cluster enrichment, log10 of p-value for cluster depletion. Dotted lines correspond to a p-value of 0.05. **F**, RNA velocity analysis shows velocity vectors from iEPs towards *Lgr5*^-^ *Lrig1*^+^ ISCs, and from the ISC clusters towards the differentiated enterocytes clusters. **G**, Subset of velocity vectors of CellTagged cells confirm transcriptional kinetics of engrafted iEPs in the direction towards intestinal stem cells.

Engrafted tissue was harvested in early stages of intestinal regeneration, with the epithelium still undergoing active repair. We chose this time point in an effort to understand the mechanism of iEP engraftment. Indeed, in line with this early regeneration period, cluster 4 likely represents cells in the early stages of engraftment and repair, characterized by expression of both intestinal and mesenchymal markers (Figs. S5D&E). Notably, Grip1, an adaptor protein implicated in maintaining the epidermal-dermal junction via the Fras1/Frem1 complex^50,51^, is among the list of marker genes for cluster 4, suggesting that cluster 4 might represent an iEP engraftment mechanism via adhesion to the basement membrane. We next focused on the proportions of fully-reprogrammed Ecad^High^ iEPs (labeled by CellTagA) and dead-end Ecad^Low^ iEPs (labeled by CellTagB) engrafting the intestine. We found that 0.687% ± 0.214% of engrafted cells were derived from reprogrammed iEPs whereas 0.413% ± 0.113% of engrafted cells were derived from dead-end iEPs (p = 0.06; Fig. S6D). This low percentage was expected given that we aimed to capture a broad range of intestinal engraftment to provide an unbiased assessment of engraftment.

In our previous study, we observed that iEPs are capable of long-term (7 weeks post-transplant), functional engraftment, where entire crypts are repopulated by iEP-derived cells^23^. At that time, we speculated that iEPs transition through an intestinal stem cell state to support long-term engraftment. Here, considering our hypothesis that fully-reprogrammed Ecad^High^ iEPs are responsible for this long-term engraftment, we performed a randomized test that we previously developed to assign statistical significance in cluster occupancy^18^. Here we applied this approach to determine whether reprogrammed and dead-end iEPs were more likely to associate with any particular cluster of intestinal cells. We did not include cluster 4 in this analysis as the colonic epithelium is in the early stages of regeneration, where we consider cells in this cluster to be superficially attached, and not all these cells will eventually integrate into the recovered epithelium. Our randomized test revealed that reprogrammed-Ecad^High^/CellTagA cells are significantly more likely to occupy cluster 0 (*Lgr5*^*-*^*Lrig1*^*+*^ ISCs, p=4.03×10^-5^) and cluster 1 (*Lgr5*^*+*^ ISCs, p=9.83×10^-3^), while CellTagged reprogrammed and dead-end populations are depleted from non-epithelial cell clusters (Fig. 3E). Together, this suggests that, Ecad^High^/CellTagA cells integrate into the regenerating epithelium via an intestinal stem cell intermediate. Expression of the ISC markers *Lgr5* and *Lrig1* in engrafted iEPs supports this observation (Fig. S6C). As reported previously, *Lrig1*^*+*^ ISCs expand and repopulate the colonic crypt upon tissue damage^27^, pointing to a potential mechanism of long-term iEP engraftment in the mouse colon.

To further investigate engraftment mechanics, we conducted RNA velocity analysis^52^ to reveal the transcriptional kinetics of engrafting iEPs. We reasoned that if these iEPs were differentiating towards intestinal lineages, then transcript kinetics from early iEP engraftment cluster, cluster 4, should show velocity vectors towards annotated intestinal clusters. Indeed, RNA velocities projected onto the UMAP clusters show cluster 4 velocities towards the main intestinal clusters (Figs. 3F, S6F&G). Specifically, velocity vectors from the subset of CellTagged cells show vectors originating from cluster 4 towards cluster 0, and from the intestinal stem cell pole of the main intestinal clusters towards the more differentiated pole of enterocytes (Fig. 3G). Taken together, here we have demonstrated the utility of CellTag Indexing to multiplex Ecad^High^ and Ecad^Low^ iEPs for transplantation into the mouse large intestine, suggesting that iEPs transition through a *Lgr5*^*+*^ and/or *Lrig1*^*+*^ stem cell state to engraft and repopulate the colonic epithelium, resolving speculation about their engraftment route. Our findings are consistent with previous reports of iEP differentiation potential and position CellTag Indexing as a powerful long-term tracking and multiplexing tool for scRNA-seq.

## DISCUSSION

Here, we introduce a broadly applicable and novel toolbox, CellTag Indexing, to label biological samples for single-cell analysis, where each sample is genetically tagged with a predefined lentiviral GFP barcode to mark its sample identity. We demonstrate that CellTag Indexing does not perturb cell physiology, and validate the utility of our multiplexing approach via species mixing, showing that it can be used to accurately multiplex samples for scRNA-seq, with subsequent demultiplexing at high-efficiency. We showcase the unique feature of this heritable labeling approach, by tracking cells in a competitive *in vivo* transplant setting, revealing reprogrammed cell potential and mechanisms of engraftment while providing internal controls to mitigate both biological and technical batch effects. CellTag multiplexing is complementary to current strategies based on transient cell surface interactions for labeling cells immediately prior to scRNA-seq, yet unique in that CellTag barcodes are stably integrated and heritable through cell division. The flexible timing of lentiviral barcode transduction, coupled with stable barcode expression, makes our system uniquely suitable for long-term tracing experiments and transplant models where temporary tags would not be retained.

CellTag Indexing offers the advantages of minimized technical variation by experimental design, the ability to multiplex biological samples for competitive transplant, broad compatibility with various cell types and single cell technologies, long-term barcode expression, streamlined workflow and library preparation, reduced sequencing cost, and straightforward demultiplexing. CellTag Indexing is designed for broad applications; its use of lentivirus as a labeling method represents a commonly used and accessible biological tool with minimal setup costs and reagent requirements. As lentivirus can transduce both dividing and non-dividing cells, CellTag barcodes can be introduced into a wide variety of cell types. In terms of estimating labeling efficiency, CellTag Indexing conveniently utilizes GFP as a barcode carrier, which can act as a visual readout for transduction efficiency. Generally, CellTag transcripts are abundantly expressed and can be optionally amplified during library preparation to further increase detection rate.

Importantly, CellTag transcripts can be recovered from the nucleus, extending this approach to single nucleus RNA-sequencing. Furthermore, cells labeled with CellTag indexes can be cultured and used in experiments prior to collection for sequencing, for example in the competitive transplant assay we demonstrated here where tagged samples act as internal controls for each other to minimize unwanted biological variation. This is complementary to existing labeling methods that utilize cell/nuclear surface chemistry or transient transfection for temporary tagging^7–12^, where the labels would be progressively lost *in vitro* and *in vivo*. Additionally, as a future application, we expect that CellTag multiplexing will be compatible with single-genome-based assays such as single-cell ATAC-seq. In summary, CellTag Indexing is a broadly-applicable tool complementary to existing methods for cell multiplexing and tracking, providing a diverse panel of experimental and analytical strategies.

## METHODS

### Cell culture

Mouse embryonic fibroblasts were derived from the C57BL/6J strain (The Jackson Laboratory 000664). HEK293T and mouse embryonic fibroblasts were cultured in Dulbecco’s Modified Eagle Medium (Gibco) supplemented with 10% Fetal Bovine Serum (Gibco), 1% penicillin/streptomycin (Gibco), and 55 μM 2-mercaptoethanol (Gibco). HAFTL pre-B cells were cultured in RPMI1640 without phenol red (Lonza) supplemented with 10% charcoal/dextran-treated FBS (Hyclone) and 55 μM 2-mercaptoethanol (Gibco)^19^.

### Generation of iEPs

Mouse embryonic fibroblasts were converted to iEPs as previously described^22,23^. Briefly, fibroblasts were prepared from E13.5 embryos, cultured on gelatin, and serially transduced every 12 hours with Hnf4α-t2a-Foxa1 retrovirus for 5 times over the course of 3 days, followed by culture on collagen in hepato-medium, which is DMEM:F-12 (Gibco) supplemented with 10% FBS, 1% penicillin/streptomycin, 55 μM 2-mercaptoethanol, 10 mM nicotinamide (Sigma-Aldrich), 100 nM dexamethasone (Sigma-Aldrich), 1 μg/mL insulin (Sigma-Aldrich), and 20 ng/ml epidermal growth factor (Sigma-Aldrich).

### CellTag barcodes

CellTag lentiviral constructs were generated by introducing an 8bp variable region into the 3’ UTR of GFP in the pSmal plasmid^53^ using a gBlock gene fragment (Integrated DNA Technologies) and megaprimer insertion (https://www.addgene.org/pooled-library/morris-lab-celltag/). Individual clones were picked and Sanger sequenced to generate predefined barcodes. The specific CellTag barcodes used in this manuscript are TGCTATAT (CellTagA), GTTGGCTA (CellTagB), AGTTTAGG (CellTagC), GGTTCACA (CellTagD), TAGAAAGC (CellTagE).

### Lenti-and retrovirus production

Lentiviruses were produced by transfecting HEK293T cells with lentiviral pSMAL vector and packing plasmids pCMV-dR8.2 dvpr (Addgene plasmid 8455) and pCMV-VSV-G (Addgene plasmid 8454) using X-tremeGENE 9 (Sigma-Aldrich). Viruses were collected 48 and 72 hours after transfection. Retroviruses were similarly produced, with retroviral pGCDNSam vector and packaging plasmid pCL-Eco (Imgenex).

### CellTag transduction

CellTag virus-containing supernatant collected from virus-producing HEK293T cells was kept at 4 °C and used within one week. Prior to transduction, protamine sulfate (Sigma-Aldrich) was added to the viral solution to a final concentration of 4 µg/ml. Cells were aspirated of media and the CellTag virus was added to the cells for a 24-hour transduction period. This transduction was repeated as needed, for a total of 48 hours for HEK293T cells and 72 hours for MEFs in the 5-tag species mixing experiment, and 72 hours for iEPs.

### Immunostaining and quantification

Transduced HEK293T and MEFs were cultured on a 4-chamber culture slide (Falcon) for 24 hr prior to fixation in 4% paraformaldehyde and staining in 300 nM DAPI in PBS. The slide was then mounted in ProLong Gold Antifade Mountant (Invitrogen). Images were acquired on a Nikon eclipse Ts2 inverted microscope. For automatic quantification, images of CellTagged HEK 293T and MEF were processed with a custom python script to count GFP positive/negative cells. The proportion of GFP positive cells was calculated from DAPI and GFP images. First, DAPI images were transformed into binary images by thresholding fluorescent signal. The threshold values were determined by the Otsu method. The binary nucleus image was processed by watershed segmentation to separate individual cell areas completely. Inappropriately sized objects were filtered to remove noise and doublet cells. The intensity of the GFP signal per individual cell area was then quantified to distinguish between GFP positive cells and negative cells. These processes were run with Python 3.6.1 and its libraries: scikit-image 0.13.0, numpy 1.12.1, matplotlib 2.0.2, seaborn 0.8.1, jupyter 1.0.0.

### Mouse model of colonic mucosal injury

Using a previously described procedure^24^, we generated colonic epithelial injury with modifications as followed: C57BL/6 mice were anesthetized with inhaled isoflurane. A custom-made syringe catheter (consisted of 3-ml syringe (BD #309657), Luer lock 26-gauge 1/2” dispensing needle (GraingerChoice #5FVG9), and polyethylene tubing (Scientific Commodities, #BB31695-PE/2) cut to approximate 5-cm in length and affixed to the needle) was used to deliver approximately 1 mL of PBS enema intraluminally via the anal canal, followed by gentle abdominal massage to promote excretion of excess fecal matter. The luminal space was then filled with 0.5 mL of 500 mM EDTA/PBS using the custom syringe catheter over the course of approximately 30 seconds. Mechanical abrasion was performed with Proxabrush cleaners (Sunstar #872FC) dipped in 500 mM EDTA/PBS, inserted approximately 1 cm into the colon, with 30 rotational movements to gently scratch the luminal surface. All animal procedures were based on animal care guidelines approved by the Institutional Animal Care and Use Committee.

### iEP characterization and transplantation

8-week iEPs were stained with mouse E-cadherin-APC antibody (10 μL per one million cells, R&D Systems, FAB7481A) and sorted on a modified Beckman Coulter MoFlo into Ecad^High^ and Ecad^Low^ populations. Sorted iEPs were plated and cultured as above. Colony formation assay was performed as previously described^18^. For colon engraftment, CellTagged Ecad^High^ and Ecad^Low^ iEPs were digested into single-cell suspensions. For each mouse, 0.5 million of Ecad^High^ iEPs (CellTagA) and 0.5 million of Ecad^Low^ iEPs (CellTagB) were pooled and resuspended in 50 μL of 10% Matrigel on ice. A total of 1 million iEPs was instilled into the colonic lumen of each mouse by using the custom syringe catheter, after which the mouse was held vertically head-down for approximately two minutes to prevent immediate excretion of the infused cell suspension.

### Single-nucleus RNA-seq procedure

Single nucleus extraction from tissue was performed as previously described^54^. Briefly, engrafted colonic tissues were finely minced with a razor then transferred to a Dounce tissue homogenizer (Kimble Chase KT885300-0002) in 2 mL of ice-cold Nuclei EZ Lysis buffer (Sigma #N-3408) supplemented with protease inhibitor (Roche #5892791001) and RNase inhibitors (Promega #N2615, Thermo Fisher Scientific #AM2696). The tissue was ground 20-30 times with the loose pestle. The homogenate was filtered through a 200-μm cell strainer (pluriSelect #43-50200) then transferred back to the Dounce homogenizer, ground with the tight pestle 10-15 times. The homogenate was incubated on ice for 5 minutes with an additional 2 mL of lysis buffer, then filtered through a 40-μm cell strainer (pluriSelect #43-50040) and centrifuged at 500 G for 5 min at 4 °C. The incubation and centrifugation steps were repeated one time, followed by resuspension Nuclei Suspension Buffer (1x PBS, 1% BSA, 0.1% RNase inhibitor) and filtering through a 5-μm cell strainer (pluriSelect #43-50005). The nuclei were then loaded onto the 10x Chromium Single Cell Platform for encapsulation and barcoding.

### scRNA-seq procedure

10x Chromium Single Cell 3’ Library & Gel Bead Kit, 10x Chromium Single Cell 3’ Chip kit, and 10x Chromium i7 Multiplex kit (10x Genomics) were used according to the manufacturer’s protocols. Libraries were quantified on the Agilent 2200 TapeStation and sequenced on Illumina HiSeq 2500.

### CellTag demultiplexing

Details of the CellTag Classifier can be found on the GitHub repository (https://github.com/morris-lab/CellTag-Classifier). Briefly, the CellTag count matrix is extracted as previously described^18^ (outlined at https://github.com/morris-lab/CloneHunter). CellTag sequences are collapsed using Starcode with the sphere clustering algorithm^55^, where CellTags with similar sequences were collapsed to the centroid CellTag. The collapsed CellTag count matrix is log-normalized, from which the most highly-expressed CellTags across cells are selected. Then, a dynamic binarization method is applied to assess the existence of each CellTag in each cell, where a ‘0’ suggests insignificant/unobservable signals and a ‘1’ indicates a significant signal. Specifically, for each CellTag, we compute the density function *D* of its expression across all cells. Then for each cell, we draw 1,000 samples from the density functions *D* and calculate the proportion *P* of samples that are greater than or equal to the expression value being tested:

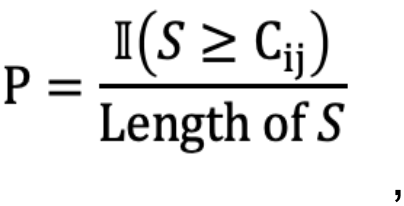

where *C*_*ij*_ = expression value of CellTag *j* in Cell *i, S* = 1,000 sample drawn from the density curve of CellTag *j, D*_*j*_. This process is iterated for at least 50 times to make sure that the samples are representative of the overall density. The cells are then classified to their corresponding CellTag based on the proportions calculated above by finding the overall minimum in each proportion matrix. The uniqueness of the minimum does not eliminate the probability for the cell to be a multiplet. Hence, for cells with a unique minimum, we examine the pair-wise differences between the minimum tag and other tags using a baseline cutoff of 0.238 learned via benchmarking and training against orthogonal 10x classification. Finally, the number of multiplets identified from our pipeline is compared to the expected number derived from 10x Genomics’ Single Cell 3’ Reagents Kit v2 User Guide Rev E (multiplet % = 0.0007589 × number of cells recovered + 0.0527214). If the number of multiplets exceeds the expected number, the optional multiplet checkpoint is implemented, where the proportion matrix is sorted such that the most likely multiplets are identified using a cutoff at the quantile of (1.5 * expected num/multiplet). The remaining cells are then classified to their singlet identities.

### scRNA-seq analysis

The Cell Ranger v.3.0.1 pipeline (https://support.10xgenomics.com/single-cell-gene-expression/software/downloads/latest) was used to process data generated using the 10x Chromium platform. For alignment of the single-nucleus RNA-seq data, a modified “pre-mRNA” mm10 reference was used to include reads aligned to introns. The R package Seurat^5^ (Version 3) was used for data processing and visualization. For the iEP dataset, we removed cells with a low number of genes detected (<200), cells with a high number of UMI detected (>100000), and cells with a high proportion of UMI counts attributed to mitochondrial genes (>20%). The filtered expression matrix was then normalized and scaled to remove unwanted sources of variation driven by number of detected UMIs and mitochondrial gene expression. Linear dimension reduction was performed, followed by canonical correlation analysis to integrate independent biological replicates, then clustering and visualization via UMAP^20^.

### Assessing cluster occupancy by randomized testing

A randomized test that we developed previously^18^ was used to identify clusters that are significantly occupied by Ecad^High^/Ecad^Low^ iEPs. In brief, we calculated the proportions of CellTagA and CellTagB cells that fall into each cluster, serving as the null percentages for the two tags. In particular, let *n* be the number of cells with a CellTag. Let *s* be the number of cells without this tag. The two were then pooled together from which we drew *n* random samples without replacement for at least *(n+s)/n* times such that every possible ending group can be captured. With each sample drawn, the occupancy of *n* sampled cells in each cluster was calculated. A background proportion distribution was then generated based on this resampling result. We then used the distributions to compute the likelihood of having the null percentage or higher. Using a p-value of <0.05, we identified the clusters that are enriched for each CellTag. This randomized test was performed using a python script. We exclude Cluster 4 in this test as it represents the early engraftment stage. Cell number tested for CellTagA = 66. Cell number tested for CellTagB = 46.

### RNA velocity analysis

RNA velocity was analyzed with Velocyto.py (version 0.17.17). The analysis was done according to the web instruction; http://velocyto.org/velocyto.py/. For the input of single-cell RNA-seq data, the output files of 10x cellranger pipeline were used. The single cell RNA-seq reads for each sample were converted into read-counts after distinguishing a spliced or unspliced transcript. This process was done with command line velocyto API and final products were saved as loom files. Next, the loom files of each scRNA-seq sample were merged into a single loom file. The merged loom file was processed with velocyto python API to create the velocyto object. Then the velocyto object was integrated with UMAP dimensional reduction data and CellTag data which were produced in the scRNA-seq analysis with Seurat and CellTag demultiplexing process. Next, the velocyto object was subjected to quality check and filtering process. Genes were filtered by the mRNA detection level (min_expr_counts=40, min_cells_express=30). After feature selection by a velocyto function, the data were normalized by total molecule count. Then velocyto object was subjected to a series of final data processing process; PCA, k-nn based imputation, velocity estimation, and shift calculation. Finally, the vectors estimated by RNA velocity was projected on the UMAP graph.

## ACKNOWLEDGEMENTS

We would like to thank members of the Morris lab and Kristen Seiler for critical discussions, John Dick for the kind gift of the pSMAL-GFP construct^53^, Genome Technology Access Center for sequencing, and Siteman Flow Cytometry core for assistance with cell sorting. This work was funded by National Institutes of Health (NIH) grants R01-GM126112, R21-HG009750, P30-DK052574; Silicon Valley Community Foundation, Chan Zuckerberg Initiative Grants HCA-A-1704-01646 and HCA2-A-1708-02799; The Children’s Discovery Institute of Washington University and St. Louis Children’s Hospital MI-II-2016-544. S.A.M. is supported by a Vallee Scholar Award; C.G.: NIH-5T32GM007200-42; K.K.: Japan Society for the Promotion of Science Postdoctoral Fellowship.

## SUPPLEMENTARY FIGURE LEGENDS

**Figure S1:**
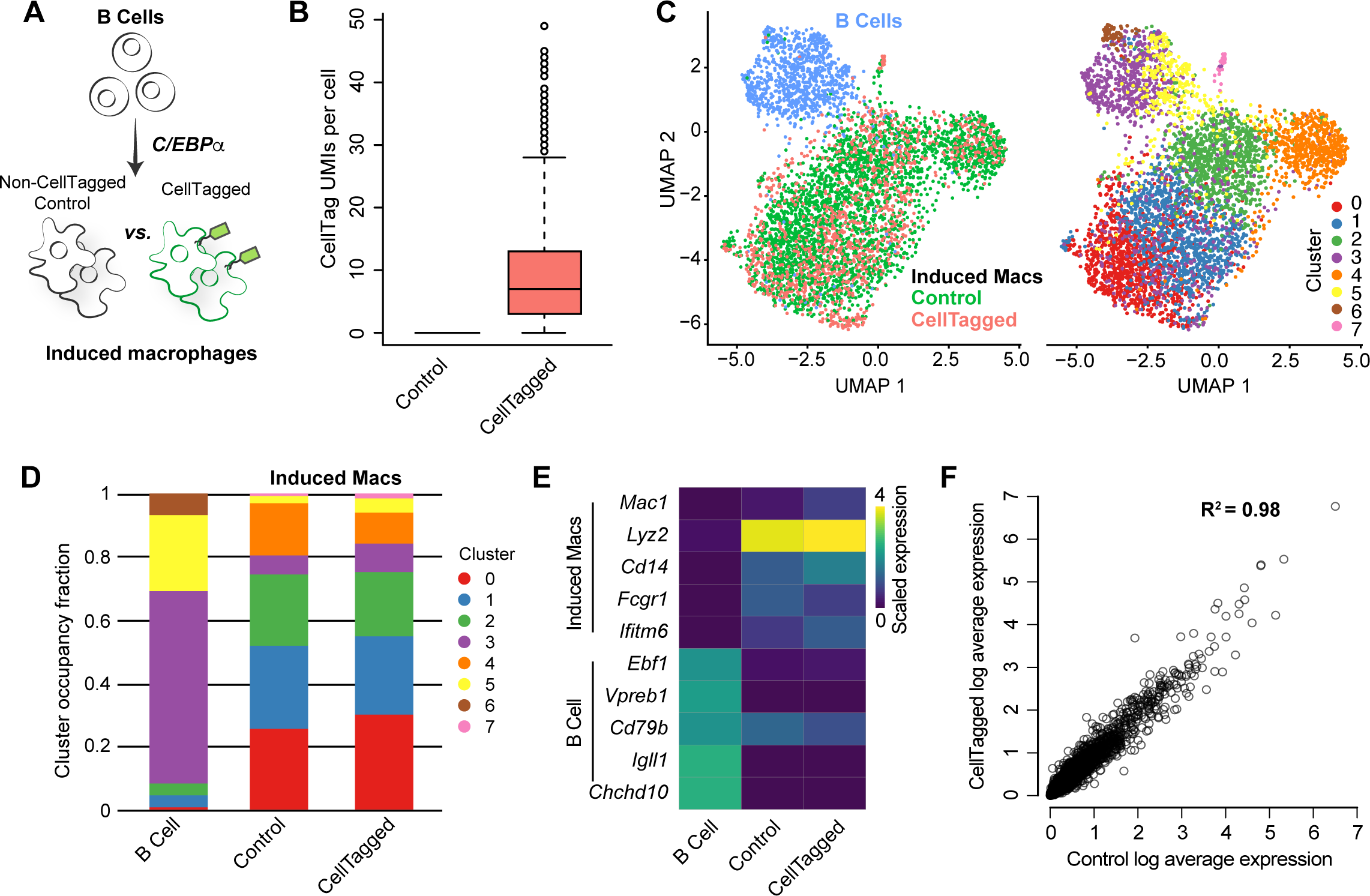
CellTag does not interfere with normal cell physiology or reprogramming. **A**, Schematic of B-cell to macrophage reprogramming and CellTagging. **B**, A median of 6 CellTag transcripts are detected per cell in CellTagged transcriptomes (CellTags were detected in all cells of this sample), while none are detected in control transcriptomes. **C**, CellTagged and control macrophages transcriptomes are interspersed with no independent clustering; both cluster separately from B cells. **D**, CellTagged and control macrophages transcriptomes share indistinguishable cluster composition, both distinct from B cell transcriptomes. **D**, CellTagged and control macrophages transcriptomes have similar macrophage marker expression and downregulate B cell marker expression. **E**, Genome-wide gene expression between CellTagged and control transcriptomes are strongly correlated with an R^2^ of 0.98.

**Figure S2:**
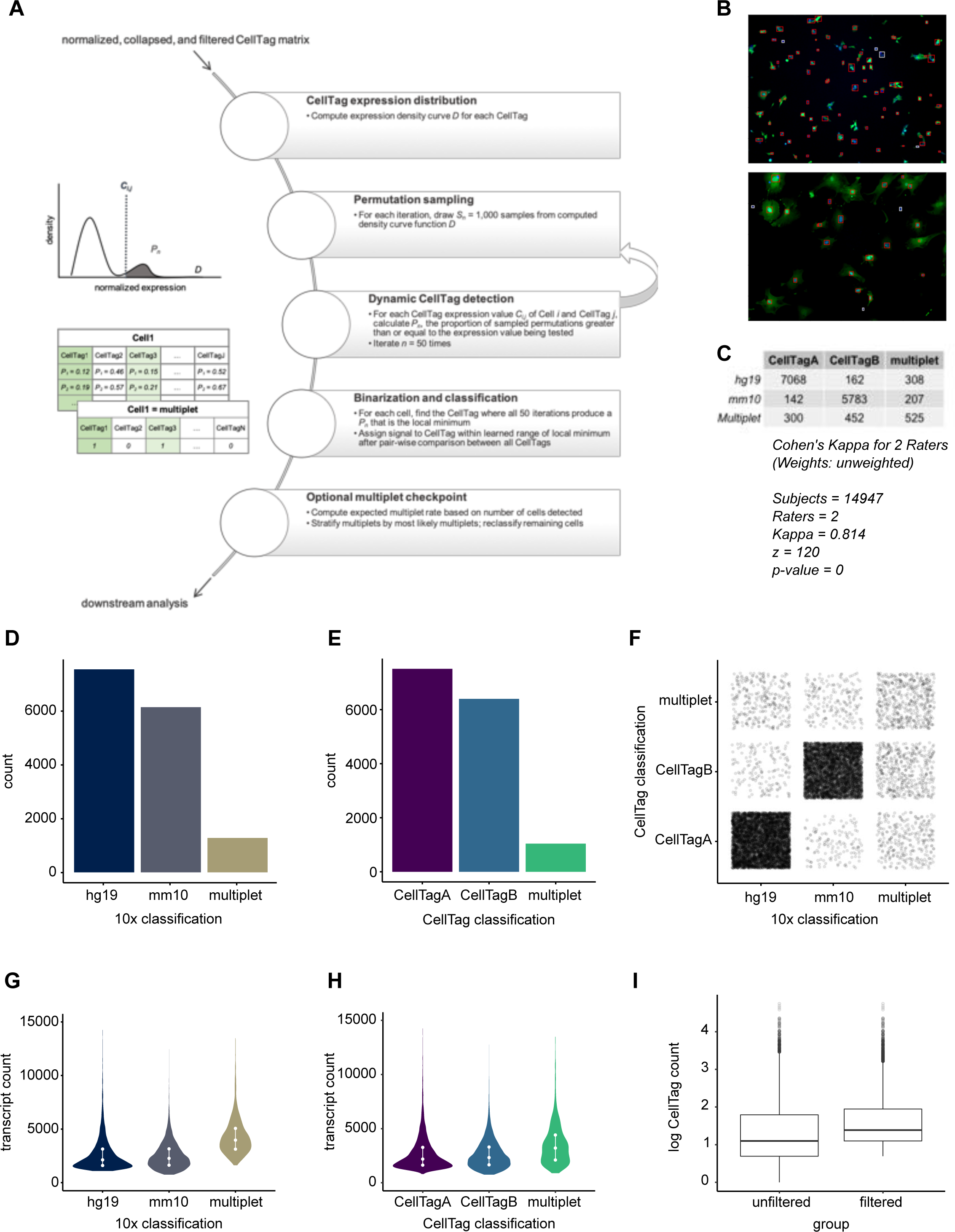
CellTag classification compared with 10x-based classification the 2-tag species mixing experiment. **A**, Schematic workflow of the dynamic binarization and classification framework. **B**, Visualization and quantification of GFP expression in DAPI-stained HEK293Ts (top, 95%) and MEFs (bottom, 88%) transduced with CellTag virus for 24-48 hours. Red box, DAPI^+^ GFP^+^; white box, DAPI^+^ GFP^-^. **C**, Table comparing CellTag and 10x-based classification, benchmarked using Cohen’s kappa as a measure of agreement. Unweighted Cohen’s Kappa = 0.814. **D**, 10x-based classification. **E**, CellTag classification. **F**, Visual comparison of CellTag and 10x-based classification. **G&H**, Total transcript count in different groups as classified by 10x-or CellTag-based classification. Dotted bars, first, median, and third quartiles. **I**, Log normalize CellTag count before and after filtering for non-determined cells.

**Figure S3:**
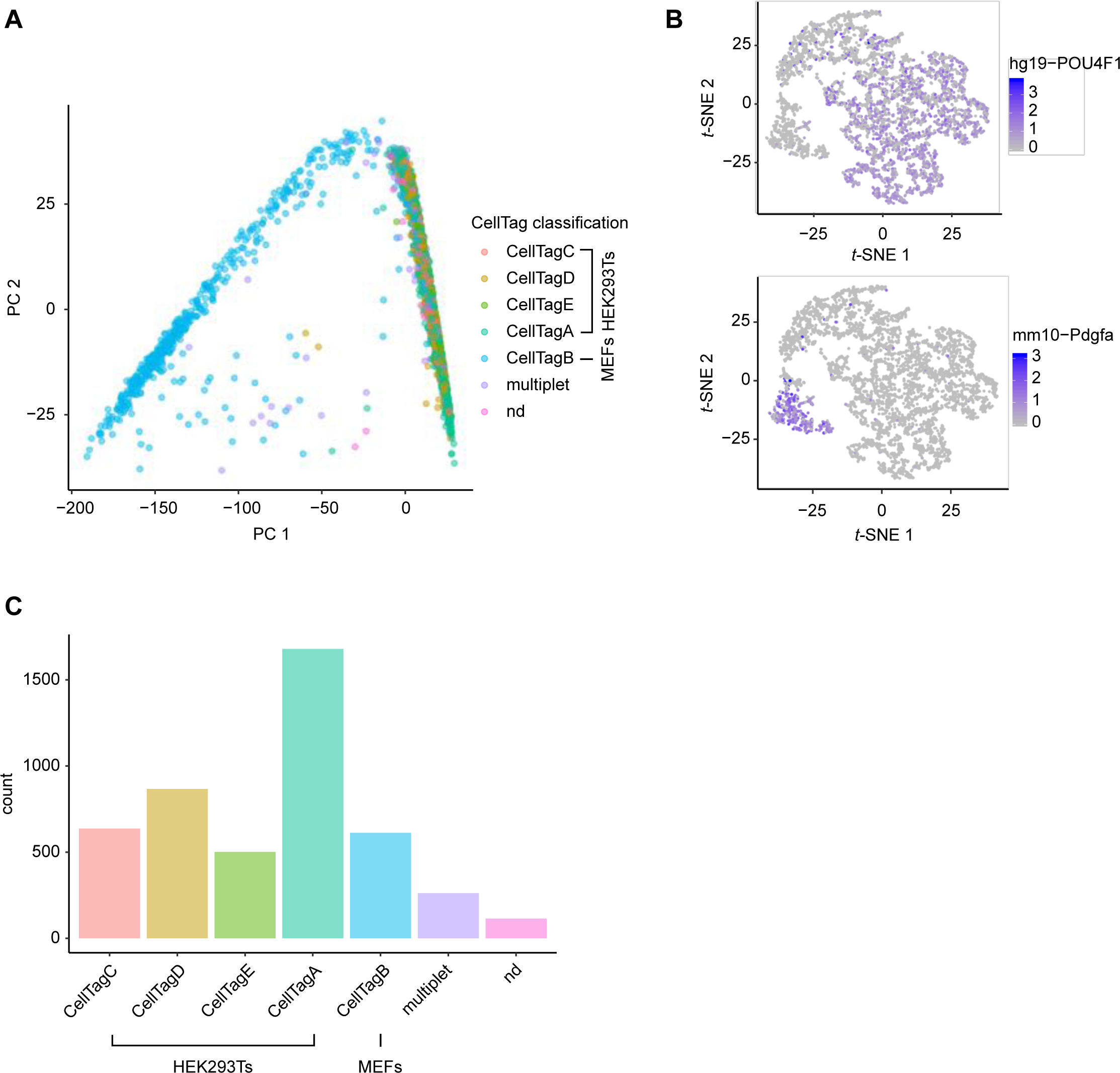
CellTag classification in 5-tag species-mixing experiment. **A**, CellTag classification visualized over transcriptomes projected onto principal component 1 (PC 1) and PC 2. **B**, Expression pattern of HEK293T marker *POU4F1* and MEF marker *Pdgfa* in mixed transcriptomes projected onto *t-*SNE. **C**, CellTag classification of the 5-tag species mixing experiment into 637 CellTagC (HEK293T), 867 CellTagD (HEK293T), 501 CellTagE (HEK293T), 1,679 CellTagA (HEK293T), 612 CellTagB (MEF), 262 multiplet, and 115 non-determined cells.

**Figure S4:**
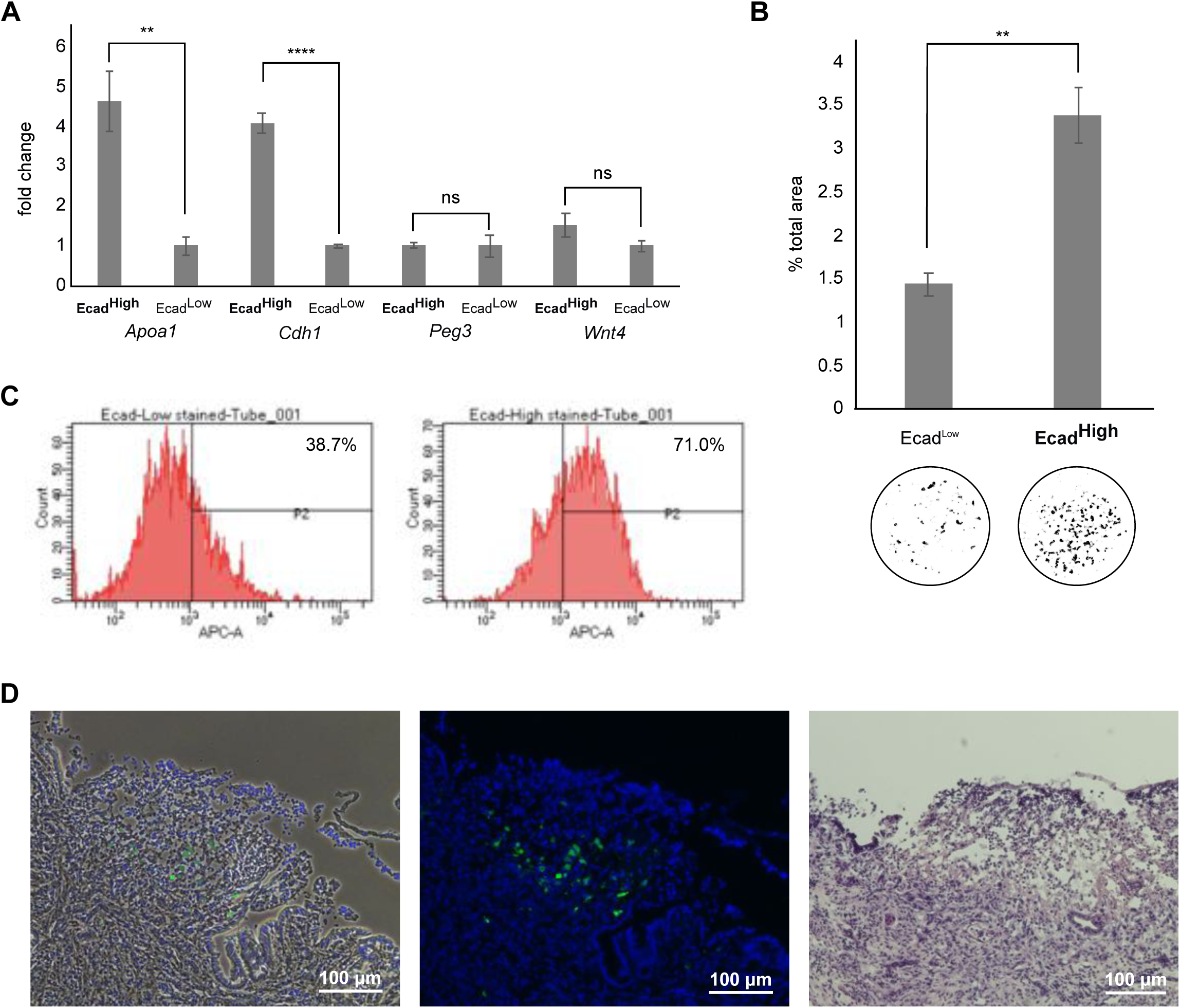
Successfully reprogrammed, Apoa1^High^Ecad^High^ iEPs can be enriched by sorting by E-cadherin expression into Ecad^High^ and Ecad^Low^ iEPs. **A**, qRT-PCR of Ecad^High^ and Ecad^Low^ iEPs show overexpression of iEP markers *Apoa1* and *Cdh1* in Ecad^High^ iEPs. **B**, Quantification of colony formation assay of Ecad^High^ and Ecad^Low^ iEPs shows that Ecad^High^ iEPs have a statistically significantly higher colony area as a proportion of total area. Bottom, threshold images of colonies. Ecad^High^ iEPs have a 1.44-fold higher colony count compared to Ecad^Low^ iEPs. **C**, Sorted Ecad^High^ and Ecad^Low^ iEPs were plated and cultured for one week. Flow cytometry analysis of cultured Ecad^High^ and Ecad^Low^ iEPs confirms that Ecad^High^ iEPs retain their Ecad^High^ phenotype. **, p < 0.01. ****, p < 0.0001. ns, non-significant. **D**, Ecad^High^ and Ecad^Low^ iEPs were labeled with CellTagA and CellTagB, respectively, and pooled in equal proportions for transplantation into the mouse colon. Left and Middle, bright field and fluorescent images of DAPI stained colon section showing aggregated GFP^+^ iEPs near the surface of the damaged epithelium. Right, H&E staining of an adjacent section showing epithelial injury and inflammation with numerous lymphocytic infiltrates. Scale bars, 100 μm.

**Figure S5:**
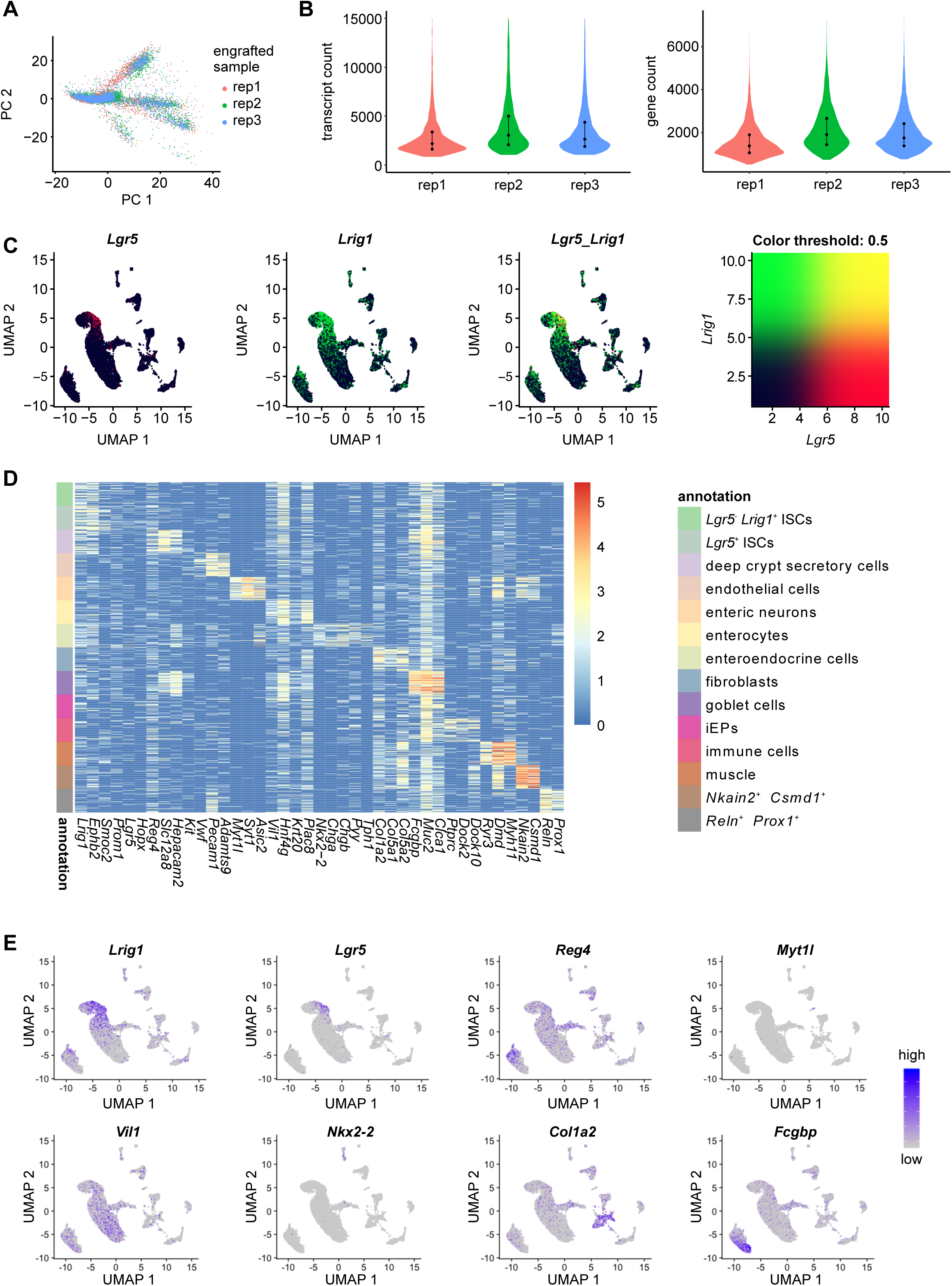
Single-nucleus RNA-seq of iEP-engrafted colon tissues reveals intestinal cell types. **A**, Visualization of three biological replicates of engrafted colon (rep1, rep2, rep3) integrated into a single dataset, projected onto PC 1 and P C2. **B**, Engrafted samples share similar levels of total numbers of transcript and gene detected per cell. **C**, ‘Blended’ feature plots of *Lgr5* and *Lrig1* expression, showing a pattern of *Lrig1* expression partially overlapping with areas with high *Lgr5* expression. **D**, Heatmap of intestinal epithelial and non-epithelial marker expression in annotated cell types (50 cells randomly sampled from each cell type). **E**, Additional feature plots of intestinal marker expression.

**Figure S6:**
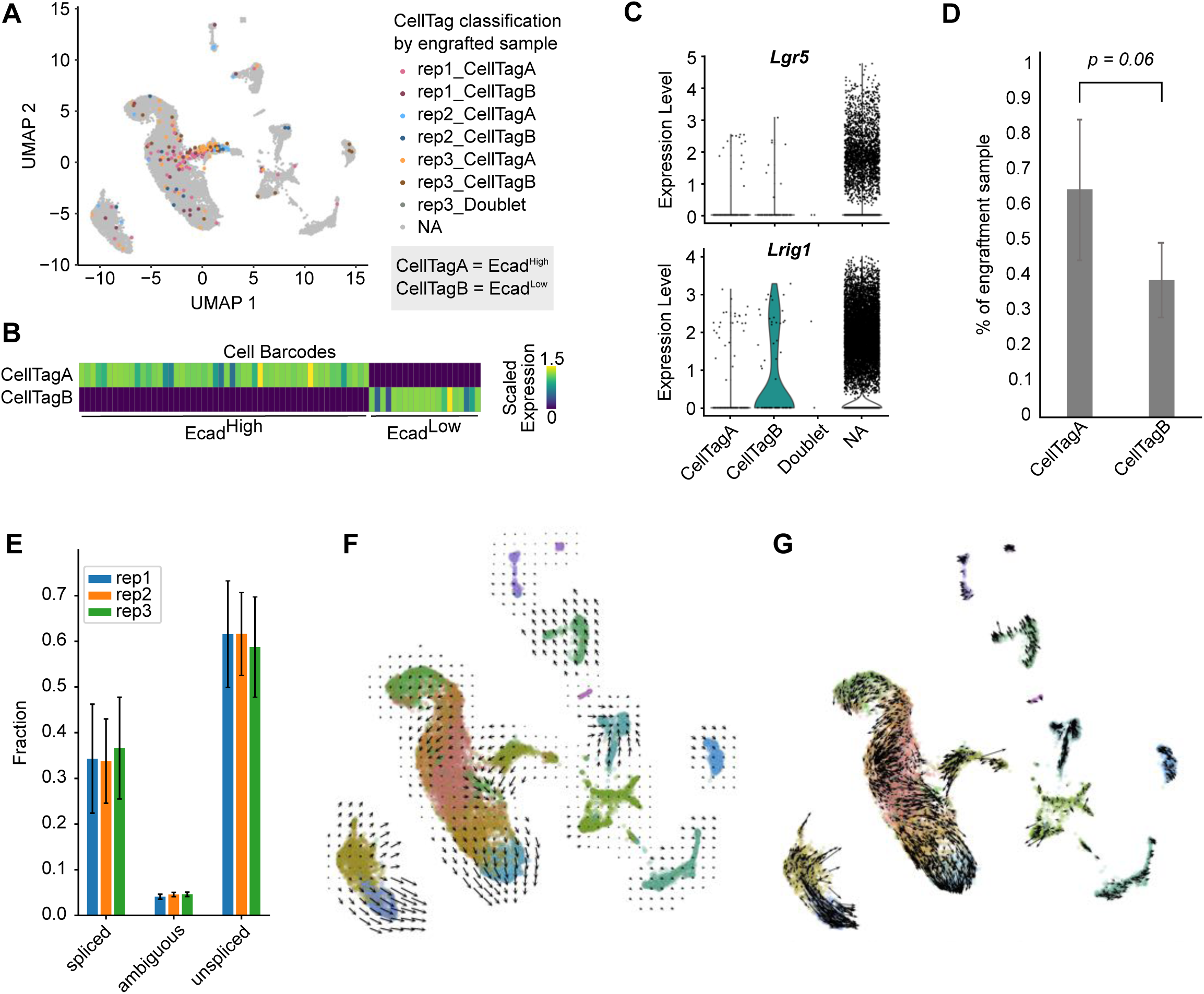
Visualization of additional CellTag and RNA velocity analysis in iEP-engrafted colon. **A**, CellTag classification shows agreement between three biological replicates of engrafted colon (rep1, rep2, rep3). **B**, Heatmap of scaled expression of CellTagA and CellTagB from rep1 shows distinct patterns of expression. **C**, 0.687% ± 0.214% of each post-engraftment sample were derived from Ecad^High^/CellTagA cells, whereas 0.413% ± 0.113% were derived from Ecad^Low^/CellTagB cells. One-sided student’s t-test, p = 0.06. **D**, *Lgr5* and *Lrig1* expression is detected in a subset of CellTagged cells. **E**, Post-engraftment samples share similar transcriptional kinetics with indistinguishable proportions of spliced and unspliced transcripts. **F**, Full vector field of RNA velocity results. **G**, Full velocity vectors of RNA velocity results.

